# A Robust High-throughput Fluorescent Polarization Assay for the Evaluation and Screening of SARS-CoV-2 Fusion Inhibitors

**DOI:** 10.1101/2021.06.17.448891

**Authors:** Xinjian Yin, Litong Chen, Siwen Yuan, Lan Liu, Zhizeng Gao

**Affiliations:** School of Marine Science, Sun Yat-sen University, Zhuhai, 519080, China; Southern Marine Science and Engineering Guangdong Laboratory (Zhuhai), Zhuhai 519080, China

## Abstract

Severe acute respiratory syndrome coronavirus-2 (SARS-CoV-2) is a serious threat to global health. One attractive antiviral target is the membrane fusion mechanism employed by the virus to gain access to the host cell. Here we report a robust protein-based fluorescent polarization assay, that mimicking the formation of the six-helix bundle (6-HB) process during the membrane fusion, for the evaluation and screening of SARS-CoV-2 fusion Inhibitors. The IC_50_ of known inhibitors, HR2P, EK1, and Salvianolic acid C (Sal C) were measured to be 6 nM, 2.5 nM, and 8.9 µM respectively. In addition, we found Sal A has a slightly lower IC_50_ (3.9 µM) than Sal C. Interesting, simple caffeic acid can also disrupt the formation of 6-HB with sub-mM concentration. A pilot high throughput screening (HTS) a small marine natural product library validates the assay with a Z’ factor close to 0.8. We envision the current assay provides a convenient way to screen SARS-CoV-2 fusion inhibitor and assess their binding affinity.

## 1. Introduction

As of Feb 2021, the severe acute respiratory syndrome coronavirus-2 (SARS-CoV-2)^[1-2]^ has infected more than 100 million people and cause two million life. Although several drugs have been “repurposed” for the treatment of SARS-CoV-2 infection,^[3-4]^ there is still a pressing need to identify novel anti-SARS-CoV-2 therapeutics.

Both SARS-CoV-1 and SARS-CoV-2 virus use their surface homotrimeric spike protein (S protein) to enter the host cells (Figure 1).^[5]^ The S protein encodes two subunits: S1 and S2. The receptor-binding domain (RBD) from the S1 subunit has a strong binding affinity with cell surface receptor, angiotensin-converting enzyme II (ACE2), and is responsible for the initial attachment.^[5]^ This binding event triggers conformational change of S2 subunit and the previously buried hydrophobic fusion peptides are exposed and insert into the host cell membrane. Subsequently, two heptad repeats (HR1 and HR2) form a thermodynamically favorably six-helix bundle (6-HB) postfusion structure to fuse the viral and cellular membranes together. ^[6]^

**Figure 1.**
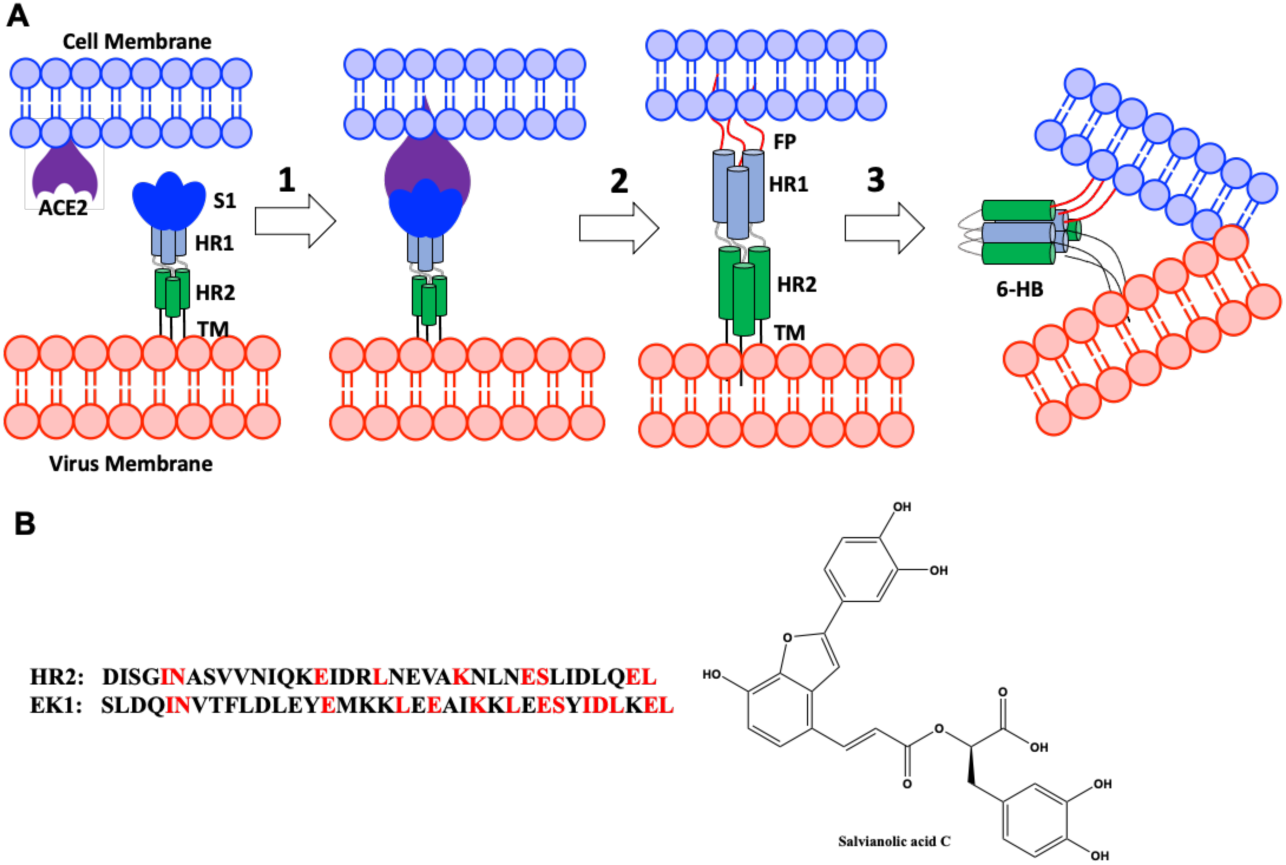
A) SARS-CoV membrane fusion mechanism. Abbreviations: FP, fusion peptide; HR, heptad repeat; TM, transmembrane anchor. B) Known SARS-CoV-2 fusion inhibitors.

The formation of 6-HB structure is a conserved fusion mechanism shared by many viruses besides coronavirus, including influenza virus, human immunodeficiency virus (HIV), and respiratory syncytial virus (hRSV).^[7-8]^ It has been shown that peptides derived from the heptad repeats region are potent fusion inhibitors, and such a peptide, Enfuvirtide, has been approved for HIV-1 treatment for almost 20 years.^[9]^ The HR2 peptide of SARS-CoV-1 and SARS-CoV-2 are 100% conserved, and also displayed potent inhibition activities in cell-based assays.^[10-13]^ Recently, the Lu group reported a novel HR2-derived peptide, EK1, which can disrupt the protein-protein interaction (PPI) during the formation of 6-HB, and inhibit multiple human coronaviruses at sub-nM concentration.^[11-12]^ Thus, fusion inhibitors could potentially be developed as broad-spectrum anti-coronavirus therapeutics.^[14]^

It is well-known that PPI is hard to disrupted by conventional small molecules.^[15]^ Natural products are privileged structures in the drug discovery process, and it might be easier to find PPI modulators from natural products ^[16]^. Recently, the Liu group identified a potent anti-SARS-CoV-2 fusion inhibitor, Salvianolic acid C (Sal-C, IC_50_ 3.85 μM) from TCM library using a cell-cell fusion assay mediated by S protein of SARS-CoV-2.^[17]^ Deconvolution of the activity by native-polyacrylamide gel electrophoresis (N-PAGE) assay indeed suggested the Sal-C inhibits the formation of 6-HB in a dose-dependent manner.

Cell-based phenotypic screening assay can be expensive and the resulting activities sometimes can be hard to deconvolute. Thus, a robust *in vitro* high-throughput evaluation and screening platform for SARS-CoV-2 fusion inhibitors is highly needed. Here we present our effort on the development of such a platform using fluorescence polarization (FP) techniques and used this platform to evaluate the binding potency of HR2, EK1, and several Sal family natural products. We found Sal-A has similar binding potency as Sal-C. In addition, we found structure as simple as caffeic acid, the precursor of Sal family natural products, can inhibit the 6-HB formation with a IC_50_ of about 638 μM. Although the potency is low, such a simple structure could potentially serve as a starting fragment for the fragment growing strategy^[18]^ to increase the potency.

## 2. Material and methods

### 2.1 Gene, Plasmid and Materials

The codon-optimized gene of 5-HB (the sequence was shown in **Table S1**) was synthesized by Tsingke Co., Ltd (Guangzhou, China) and inserted into plasmid pET-44b(+) and pET-28a(+), respectively. The pCold-NusA plasmid was maintained in our laboratory.

The HR1P (CANQENSAIGKIQDSLSSTASALGKLQDVVNQNAQALNTLVKQ) and HR2P (ELGDISGINASVVNIQKEIDRLNEVAKNLNESLIDLQELC) peptides were ordered from APeptide Co., Ltd (Shanghai, China). EK1 (SLDQINVTFLDLEYEMKKLEEAIKKLEESYIDLKEL) peptide was a gift from Prof. Lu Lu (School of Basic Medical Sciences, Fudan-Jinbo Joint Research Center, Fudan University, Shanghai 200032, China). Salvianolic acid A, Salvianolic acid B, Lithospermic acid, Rosmarinic acid, and Caffeic acid were purchased from Aladdin Reagents Co. Ltd (Shanghai, China). Salvianolic acid C was purchased from MACKLIN Reagent Co. Ltd (Shanghai, China). All other chemicals and reagents were chemically pure and were obtained commercially.

### 2.2 Protein expression and purification

The plasmids pET-44b-NusA-5HB, pET-28a-5HB, and pCold-NusA containing the gene encoding NusA-5HB, 5HB and NusA (with N-terminal His-tag) were transformed into competent *E. coli* BL21 (DE3) cells (Invitrogen) respectively. The recombinant *E. coli* was cultivated in LB medium with antibiotic (50 mg/L Kan for pET-28a-5HB; 100 mg/L Amp for pET-44b-NusA-5HB and pCold-NusA) at 37 °C and rotary shaking at 200 rpm. For protein expression, cells were induced with isopropyl-β-D-thiogalactopyranoside (IPTG) at a final concentration of 0.5 mM until OD_600_ reached 0.6-0.8, and then further cultured at 15 °C for 24 h.

The collected cells were washed and resuspended in buffer A (20 mM sodium phosphate, pH 7.4, containing 500 mM NaCl and 20 mM imidazole). Resuspended cells were disrupted by ultrasonication in an ice bath, followed by centrifugation at 12000 × g for 30 min to remove cell debris. The supernatant was loaded onto a Ni-NTA column (Thermo Scientific, USA) pre-equilibrated with buffer A, and the proteins were eluted by an increasing gradient of imidazole (from 50 to 250 mM). The purities of the collected fractions were analyzed by SDS-PAGE. Fractions containing the pure target protein were gathered and then desalted by ultrafiltration. The purified proteins were concentrated and stored in 20% (v/v) glycerol at −80 °C until further use. The expression and purification of the proteins were analyzed by sodium dodecyl sulfate-polyacrylamide gel electrophoresis (SDS-PAGE, 4∼20%) (Figure S1). Protein concentrations were determined using a Bradford protein assay kit (Quick Start™, Bio-Rad, USA).

### 2.3 HR2P-FL preparation

Dissolving HR2 with 20 mM pH 7.5 phosphate buffer (5 mM EDTA) to a concentration of 100 μM. A 25-fold molar excess of fluorescein-5-maleimide (2.5 mM) was added and then incubated at 25 °C for 2 h. After incubation, nonreacted fluorescein was removed by ultrafiltration using a 3 KDa molecular-weight cutoff, and HR2P-FL was then purified by HPLC. The purified HR2P-FL was stored from light in single-use aliquots at -20 °C.

### 2.4 Fluorescence Native polyacrylamide gel electrophoresis (N-PAGE)

10 µM of HR2P-FL was incubated with 10 µM of HR1P, 5HB or NusA-5HB in pH 7.5 20 mM PBS at 37 °C for 30 min. After incubation, the sample was mixed with Tris-glycine native sample buffer (Invitrogen, Carlsbad, CA) at a ratio of 1:1 and was then loaded to 4%∼20% pre-cast gel (20 μL each well). Gel electrophoresis was carried out with 100 V constant voltages at room temperature for 1.5 h. The gel was imaged with a Tanon 1220 Gel Imaging System (Shanghai, China)_º_

### 2.5 FP Assay

All reported concentrations represent the final assay conditions unless otherwise specified. Fluorescent polarization measurements were performed on a Spark® multimode microplate reader (Tecan, Switzerland), with an excitation filter at 485 nm, and an emission filter at 535 nm. The G-Factor was set at 1.06 (Note: The calibration is performed by measuring a 1 nM fluorescein solution at room temperature and adjusting the G-factor to achieve a value of 27 mP; different microplate readers might need a different setting)^[19]^. Peptides were directly dissolved in water to make a 500 µM stock solution and then stored at -80 °C.

#### 2.5.1 Binding assay of HR2-FL with NusA-5HB/5HB

Assays were performed manually in black 96 microplates (Cat. No. 3694, Corning, USA). Each reaction (100 µL) contained 10 nM HR2P-FL and 1 µM of NusA-5HB, 5HB or NusA in FP buffer (25 mM pH 7.5 PBS, 0.025% NP-40). The FP values were measured after 1 h incubation at 37 °C 200 rpm. All experiments were performed in duplicate.

#### 2.5.2 Saturation binding FP measurement

Assays were performed manually in black 96 microplates (Cat. No. 3694, Corning, USA). Each well (100 µL) contained 5 nM HR2P-FL and increasing concentrations (0∼128 nM) of NusA-5HB in FP buffer (25 mM pH 7.5 PBS, 0.025% NP-40). The FP values were measured after 1 h incubation at 37 °C 200 rpm. The binding fraction (*f*) of HR2-FL with different NusA-5HB concentrations was calculated using the **Equation 1**,**2**.^[20]^ The obtained *f* values were analyzed and plotted using GraphPad Prism 8 to calculate the binding dissociation constant (*K*_d_) by fitting the experimental data using a one-site specific binding model. All experiments were performed in duplicate.

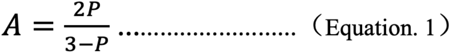

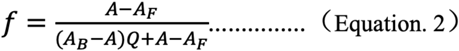

*P* is the fluorescence polarization values, *A* is the fluorescence anisotropy values; *A*_*B*_ and *A*_*F*_ denote fluorescence anisotropies of fully bound and free HR2P-FL, respectively; Q is the ratio of fluorescence intensities of bound and free species measured under the same experimental conditions. In our assay, Q is found to be close to 1.

#### 2.5.3 Competitive FP assays of peptides

Assays were performed manually in black 96 microplates (Cat. No. 3694, Corning, USA) contained 8 nM NusA-5HB and increasing concentrations of peptides (0, 0.31, 0.625, 1.25, 2.5, 5, 10, 20, 40, 80,160, 320 nM) in FP buffer (25 mM pH 7.5 PBS, 0.025% NP-40) in a final volume of 90 μL. After 1 h incubation at 37 °C 200 rpm, 10 μL HR2P-FL (5 nM) was added and additional 1 h incubation was followed at 37 °C 200 rpm. The FP signal was measured. The binding fraction (*f*) of HR2-FL with NusA-5HB at different concentrations of peptide was calculated using the Equation 1,2. The obtained *f* values were plotted using GraphPad Prism 8 to calculate the IC_50_. All experiments were performed in duplicate.

#### 2.5.4 Competitive FP assays of small molecules

Assays were performed manually in black 96 microplates (Cat. No. 3694, Corning, USA) contained 8 nM NusA-5HB and increasing concentrations of small molecules (0, 0.001, 0.01, 0.1, 1, 10, 100, 1000 μM) in FP buffer (25 mM pH 7.5 PBS, 0.025% NP-40, 4% DMSO) in a final volume of 90 μL. After 1 h incubation at 37 °C 200 rpm, 10 μL of HR2-FL (5 nM) was added and additional 1 h incubation was followed at 37 °C 200 rpm. In order to eliminate the fluorescence interference of small molecules, fluorescence intensities parallel (*I*_*=*_), and perpendicular (*I*_*⊥*_) to the plane of exciting light were measured after incubation and the real FP values were calculated using the Equation 3. The binding fractions (*f*) of HR2P-FL with NusA-5HB at different concentrations of small molecules were calculated using the Equation 1,2. The obtained *f* values were plotted using GraphPad Prism 8 to calculate the IC_50_. All experiments were performed in duplicate.

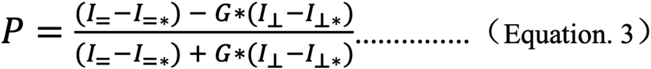

*I*_*=\**_ and *I*_*⊥\**_ are fluorescence intensities of same concentration small molecules parallel, and perpendicular to the plane of exciting light. G is the G-Factor (1 nM fluorescein solution at room temperature and adjusting the G-factor to achieve a value of 27 mP).

### 2.6 Molecular Docking

The solved post-fusion core of SARS-CoV-2 6-HB was selected as the docking structure (PDB code: 6LXT)^[11]^. In this structure, one of the HR2 chains was removed and the channel occupied by this HR2 was defined as the docking grid. Ligands were prepared using MGLTools 1.5.4 and docking was performed with auto-dock vina. The molecular docking poses were ranked according to calculated binding affinity and the top one was selected for further analysis. PyMOL (Version 1.5, Schrö dinger, LLC) was used to prepare figures of the complexes.

### 2.7 Screening the marine natural product library

The marine natural product library (10 mM dissolved in 100% DMSO) in 96-well format was maintained in our laboratory. A daughter library (100 times dilution) was created by transferring 1 µL of the stocks into 99 µL of 40% DMSO. To black 96-well microtiter plates were added 80 µL of a premix containing 8 nM NusA-5HB, buffer (25 mM pH 7.5 PBS, 0.025% NP-40). 10 µL daughter library solution was added (10 µM) and incubated at 37 °C for 1 h. Then, 10 µL of HR2P-FL (5 nM) was added and additional 1 h incubation was followed at 37 °C 200 rpm. FP signal was measured after incubation. (Note: All concentrations represent the final assay conditions)

For quality assessing of the FP assay, positive and negative controls were set in the same 96-well microtiter plates and subjected to identical conditions described in the above screening procedure. Positive controls refer to the reactions in the absence of inhibitors while negative controls refer to the reactions in the absence of NusA-5HB. Z’ score was calculated according to the equation as follows ^[21]^:

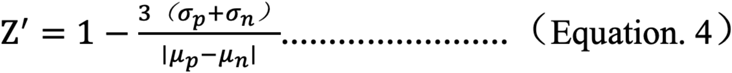

The and are the standard deviations of the FP signals of positive and negative controls; and are the mean of FP signals of positive and negative controls.

## 3. Results

### 3.1 FP-based assay for fusion inhibitors

FP is a powerful technique for the investigation of ligan-receptor interactions in homogenous solutions, and has been widely used in the drug discovery process.^[19, 22]^ The basic concept is to link a fluorophore to the ligand and the fluorophore-ligand conjugate has a low molecular weight, thus freely rotating in the solution to give a low FP signal. Upon the addition of the receptor (typical an enzyme with higher molecular weight), the binding between receptor-ligand would non-covalently link the fluorophore to a large object, and the rotation is restricted and a high FP signal is obtained.

The Kim group previously design a small protein, 5-HB, by connecting the five of the 6-HB of HIV-1 with short linkers.^[23]^ The resulting 5-HB have a strong binding affinity with the HR2 region of HIV-1 and displaced a strong fusion inhibition activity. Subsequently, such a 5-HB construct was used in an *in vitro* FP assay for screening HIV fusion inhibitors.^[24]^ Inspired by those results, we developed an FP-based assay for SARS-CoV-2 fusion inhibitors as shown in Figure 2. An expression plasmid with short linkers to link three HR1 and two HR2 of SARS-CoV-2 was constructed. The resulting 5-HB construct would be expected to have a high binding affinity with fluorescence labeled HR2 peptides (HR2P) to increase the FP signal. An extra cysteine is included in the synthetic HR2P (there is no cysteine in the sequence) to facilitate the labeling with fluorescein-5-maleimide to get HR2P-FL.

**Figure 2.**
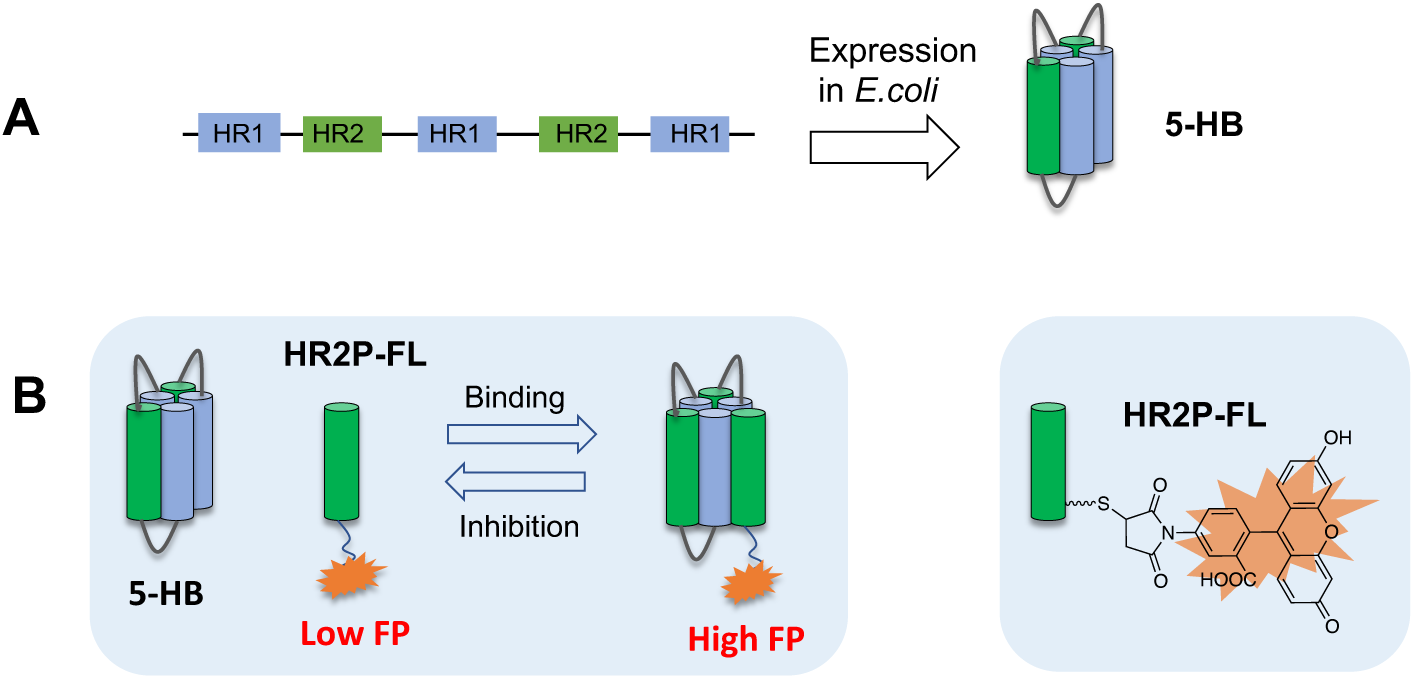
The FP-based assay for SARS-CoV-2 fusion inhibitor. A) 5HB expression construct. B) FP assay. Upon 5HB binding to HR2P-FL, the FP reading of HR2P-FL increases. If inhibitors can disrupt the binding between 5-HB and HR2P-FL, the FP reading decreases.

We found 5HB is mostly expressed in *E. coli*. exclusion body and the yield of soluble 5HB fraction is very low. We envision that a solubilization protein tag would aid the expression. Here we chose the *E. coli*-derived NusA tag (55 kDa)^[25-26]^ as it has high intrinsic solubility and its large size can potentially increase the FP signal. Indeed, NusA tag has been employed to help the expression of HIV-5HB.^[27]^ We are glad to found that NusA-5HB construct gave a very high expression yield, and about 30 mg-50 mg/L of soluble NusA-5HB can be obtained after purification.

### 3.2 Validation of the FP-based assay

With the HR2P-FL, NusA-5HB, and 5HB in hand, we first mixed 10 nM of HR2P-FL and excess of NusA-5HB and 5HB (1 μM). After incubation for 1 h, the FP reading was measured by a plate reader. HR2P-FL gave a relatively low FP reading of 50 mP, while NusA-5HB and 5HB raise the FP reading to 220 and 230 mP, respectively (Figure 3A). Although NusA-5HB (100 kDa) is significantly larger than 5HB (35 kDa), comparable FP readings were obtained. This may be because the internal linker between NusA and 5HB is flexible and the NusA can not restrict the rotation of 5HB.^[28-29]^ To rule out that NusA itself would bind to HR2P-FL to increase the FP reading, we also expressed the NusA protein, and we found NusA did not change the FP reading, suggested NusA does not bind to HR2P-FL.

**Figure 3.**
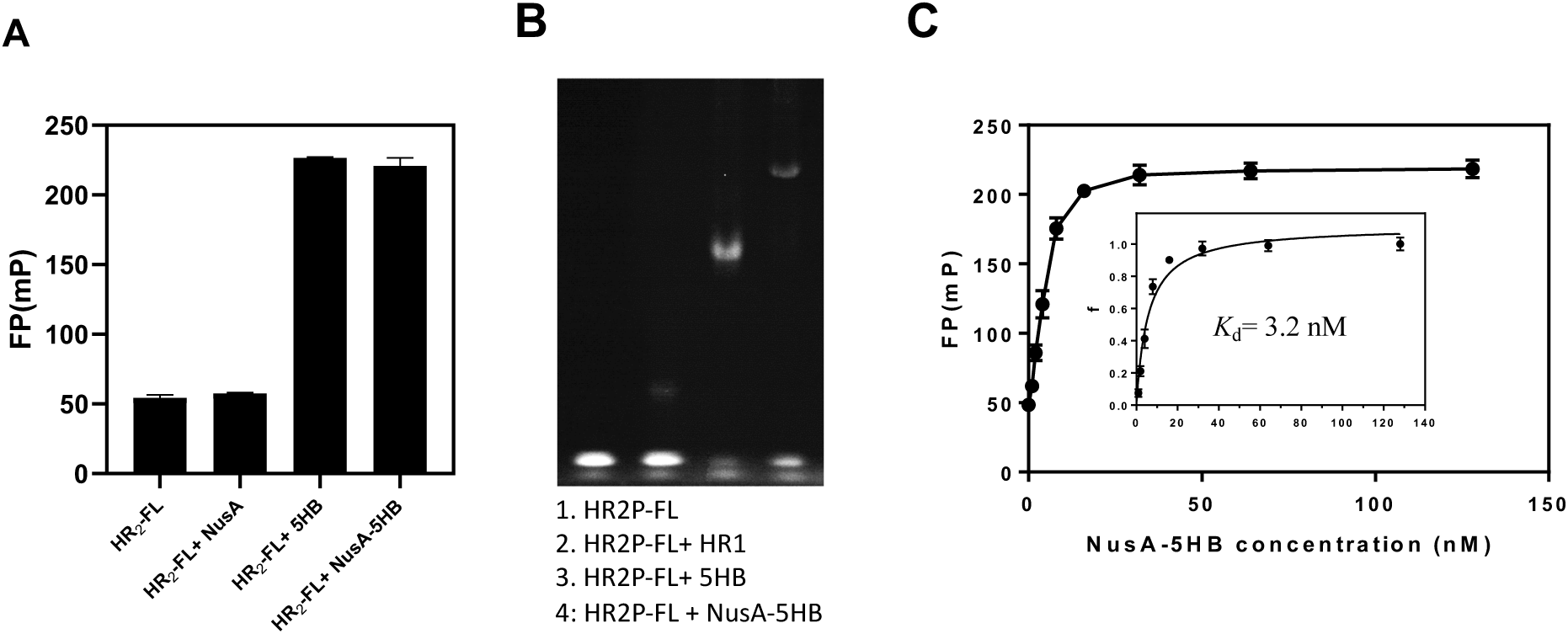
Validation of the FP assay. A) FP readings of 10 nM of HR2P-FL; 10 nM of HR2P-FL with 1 μM of NusA, 5HB, or NusA-5HB. B) Determination of the six-helix bundle formation between HR2-FL and 5HB/NusA-5HB by FN-PAGE. Lane 1: 10 μM HR2P-FL; Lane 2: 10 μM HR2P-FL+10 μM HR1; Lane 3: 10 μM HR2P-FL+10 μM 5HB; Lane 4: 10 μM HR2P-FL+10 μM Nusa-5HB. C) Saturation binding FP experiments. The FP readings were converted to fraction bounded (*f*), and then plotted against the NusA-5HB concentration to get *K*_d_ = 3.2 nM.

To further validate the binding between HR2P-FL and 5HB, we performed native-polyacrylamide gel electrophoresis (N-PAGE) analysis.^[12]^ As shown in Figure 3B, we found HR2P-FL and HR1P only formed a small amount of 6-HB, while HR2P-FL can form 6-HB effectively with either 5HB and NusA-5HB. Since both experiments suggest HR2P-FL could bind to NusA5-HB to form NusA-6HB, we subsequently focused on the assay optimization with NusA-5HB.

### 3.3 Saturation binding FP experiments

To develop a successful FP-based assay, the *K*_d_ between fluorescent ligand and receptor need to be determined by saturation binding experiment.^[20, 30]^ The concentrations of fluorescent ligand and receptor in the subsequence competitive FP assay need to be adjusted according to the saturation binding experiment:the fluorescent ligand concentration needs to be less than 2 *K*_d_ to avoid stoichiometric titration; while the concentration of receptor needs to be adjusted so that the initial fraction of fluorescent ligand bound to receptor (*f*_0_) is between 0.5 to 0.8.

To determine the binding constant between NusA-5HB and HR2P-FL, increasing concentrations of NusA-5HB were added to the solution containing 5 nM HR2P-FL, and after incubate for 1 h, FP signals were measured ((Figure 3C)). As FP signals are not linear with the binding strength,^[20]^ the anisotropy data were used to calculate the fraction bounded ratio (*f*), which were then plotted against the NusA-5HB concentrations to get the *K*_d_ = 3.2 nM.

### 3.4 Competitive FP assays with HR2P and EK1

With the saturation binding experiment data in hand, we set the HR2P-FL and NusA-5HB concentrations to be 5 nM (less than 2 *K*_d_) and 8 nM (*f*_0_ is about 0.7) respectively in the competitive FP assays. As shown in Figure 4, IC_50_ of HR2P and EK1 is 6.1 and 2.5 nM respectively, suggesting that EK1 has a slightly higher binding toward 5HB. However, it has been shown that HR2 has better activity than EK1 in cell-based pseudovirus infection assay (0.98 μM vs 2.38 μM). It is possible that the cell-based assay does not reflect the true binding affinity as peptides might have different cell-penetration abilities. Indeed, the Lu group recently reported that cholesterol or palmitic acid conjugated EK1 have much better *in vivo* activity than EK1, and the IC_50_ of both labeled peptide are in nM range.

**Figure 4.**
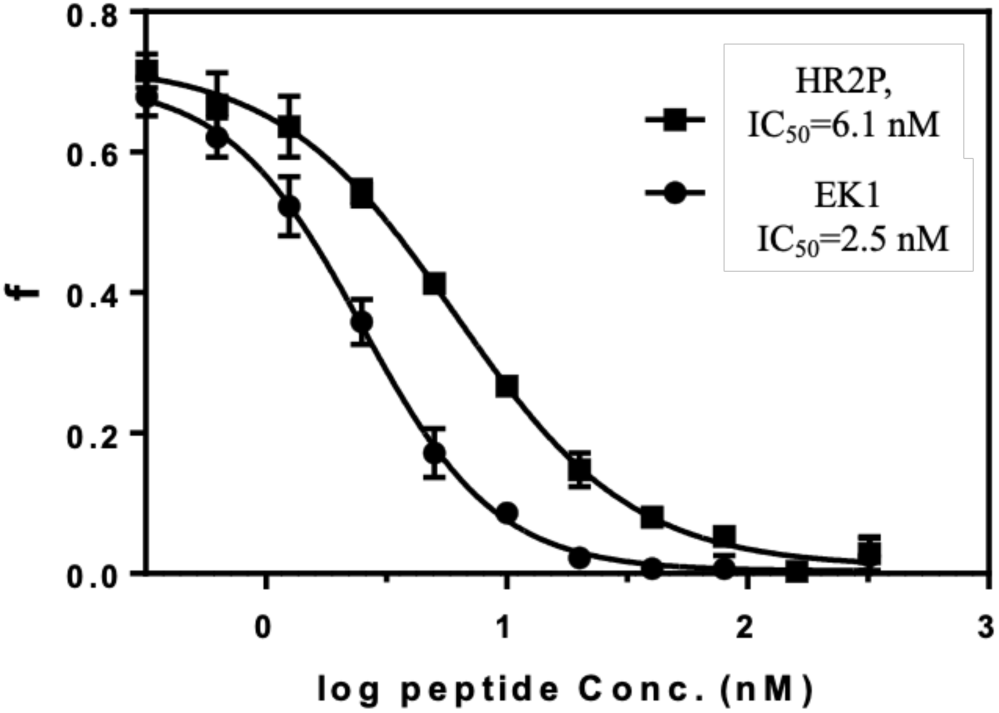
Competitive FP assays with HR2P and EK1. The FP readings were converted to fraction bounded (*f*), and then plotted against the HR2P and EK1 concentration to get IC_50_ = 6.1 nM for HR2P and IC_50_ = 2.5 nM for EK1, respectively.

### 3.5 Evaluation of Sal family natural products with FP assay

The Liu group reported that nonpeptic natural product Sal-C is a potent SARS-CoV-2 fusion inhibitor with a IC_50_ of 3.85 µM with an ACE2-expressing HEK293T cell, and a EC_50_ of 3.41 µM with native SARS-CoV-2 virus.^[17]^ The mode of action of Sal-C was elucidated by N-PAGE, and it was observed that Sal C inhibits the formation of 6-HB between HR1P and HR2P in a dose - dependent manner. However, the binding affinity between Sal-C and SARS-CoV-S2, SARS-CoV-2-S2 (405 µM and 284.3 µM respectively) is relatively low, possibly because S2 unit is not in the prefusion state. Our current assay has the potential to evaluate the true binding strength between Sal-C and spike protein in its prefusion state. In a competitive bind experiment, we obtained the IC_50_ of Sal C to be 8.9 µM (Figure 5), comparable to that determined in the cell-based assay, suggest that Sal C is possibly interacts at interface of 5HB and HR2P. This result strongly suggested that our assay can be used to evaluate the potency of SARS-CoV-2 fusion inhibitors.

**Figure 5.**
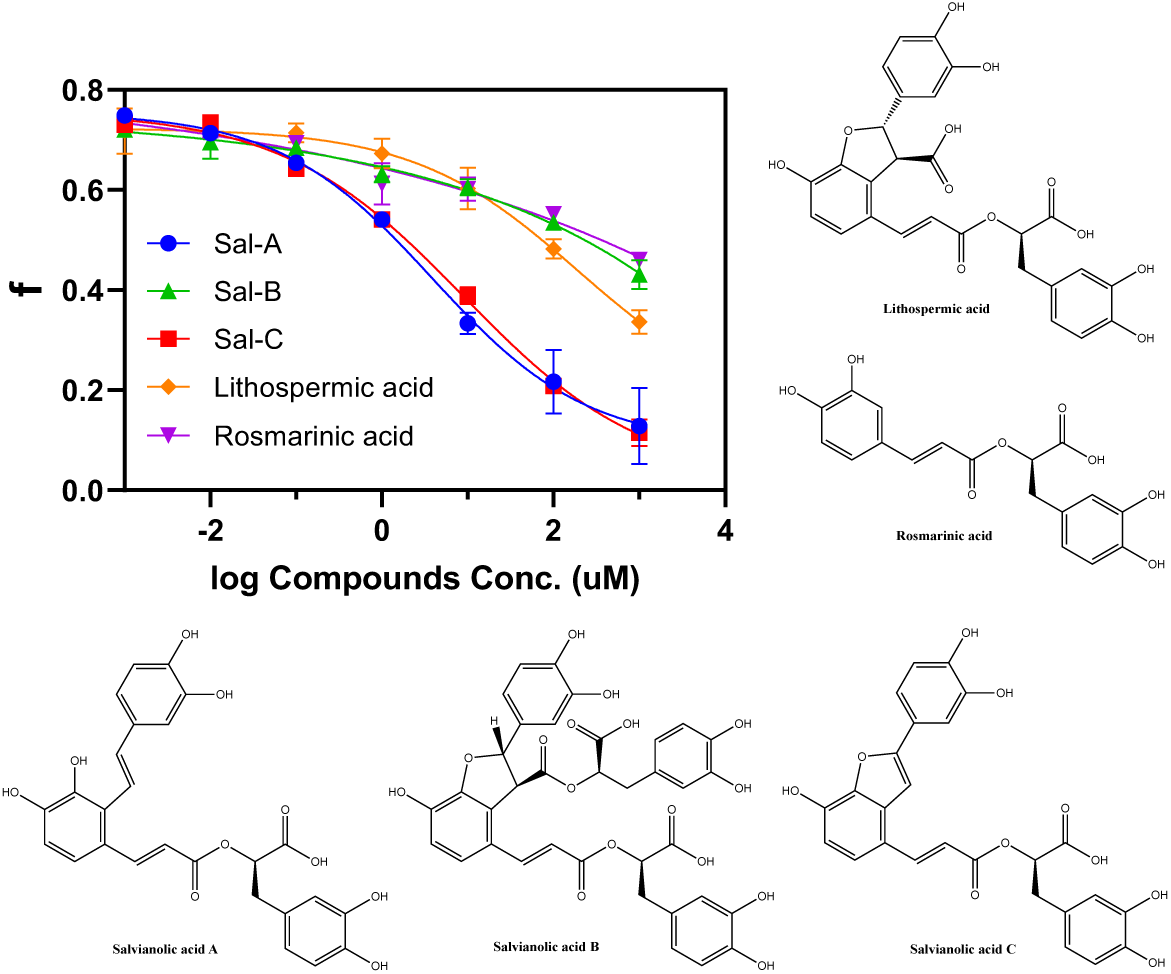
Evaluation of Sal family natural products. The FP readings were converted to fraction bounded (*f*), and then plotted against the respective natural product concentrations.

With these encouraging results in hand, we tested several Sal C analogs that are commercially available: Sal A, Sal B, lithospermic acid, rosmarinic acid. The IC_50_ of Sal A is about 3.9 µM, slightly better that Sal C, suggesting the furan ring of Sal C is not necessary. The other three natural products all have very week activity, and because of solubility issues, the IC_50_ values can not be obtained. Those results suggest although the furan ring is not necessary, the double bond of furan ring is crucial for the activities.

Since Sal family natural products are oligomers of caffeic acid, we are curious that caffeic acid can also bind to 5-HB. Since caffeic acid have higher solubility, we successfully obtained a IC_50_ of 638 µM (Figure 6A). PPI does not have a binding pocket, and usually it is hard to find binders with simple structures. It is pretty remarkable a simple structure such as caffeic acid could disrupt the interaction between 5HB and HR2P in less than mM concentration. We then performed docking studies with the solved 6-HB crystal structure (PDB code:6LXT)^[11]^, and it suggests that the carboxyl group of caffeic acid forms hydrogen bonding with Lys947, and para phenolic OH might interact with Ser-940 (Figure 6B). We envision that caffeic acid could potentially serve as a starting fragment in fragment-based drug discovery to increase the binding with 5HB.

**Figure 6.**
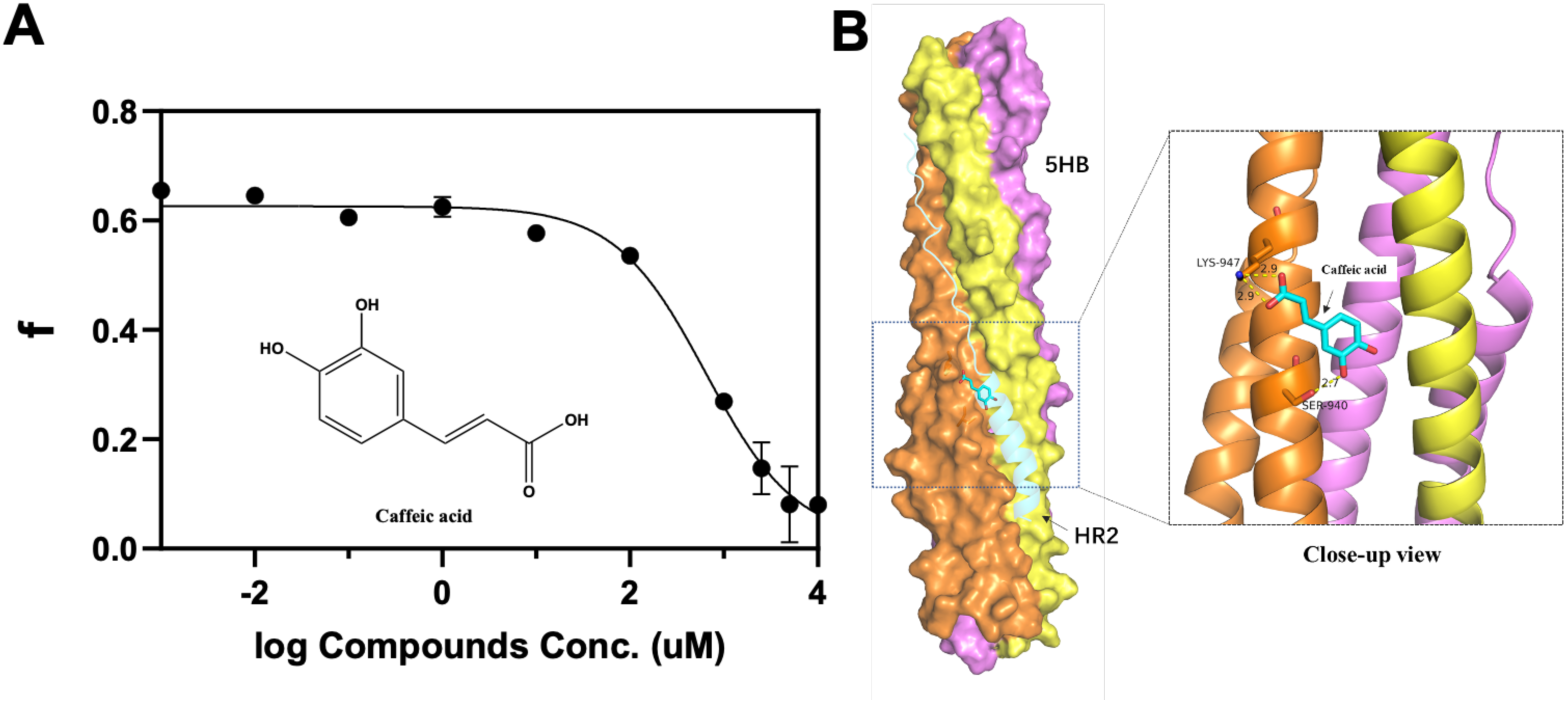
Evaluation of the caffeic acid activity. A) IC_50_ of caffeic acid. The FP readings were converted to fraction bounded (*f*), and then plotted against the respective natural product concentrations. B) Docking structure between caffeic acid and 6-HB without one of HR2 (PDB code:6LXT).

### 3.6 High-throughput screening (HTS) of marine natural product library

As a marine natural product research group, we have a long-term interest to find bioactive molecules from the marine environment, and indeed we have previously successfully identified several potent inhibitors from marine natural product libraries with FP-based assays^[31-32]^. Therefore, we performed a pilot HTS with a small marine natural product library (about 300 compounds) at 10 µM (Figure 7A). Unfortunately, we did not find any active natural products from this small library after ruling out the fluorescent interfering compounds. However, we found the assay itself is very robust with a Z’ at 0.77 (Figure 7B), which suggested that this assay can be applied to screen large-size commercial compound libraries, which unfortunately we currently do not have assess to.

**Figure 7.**
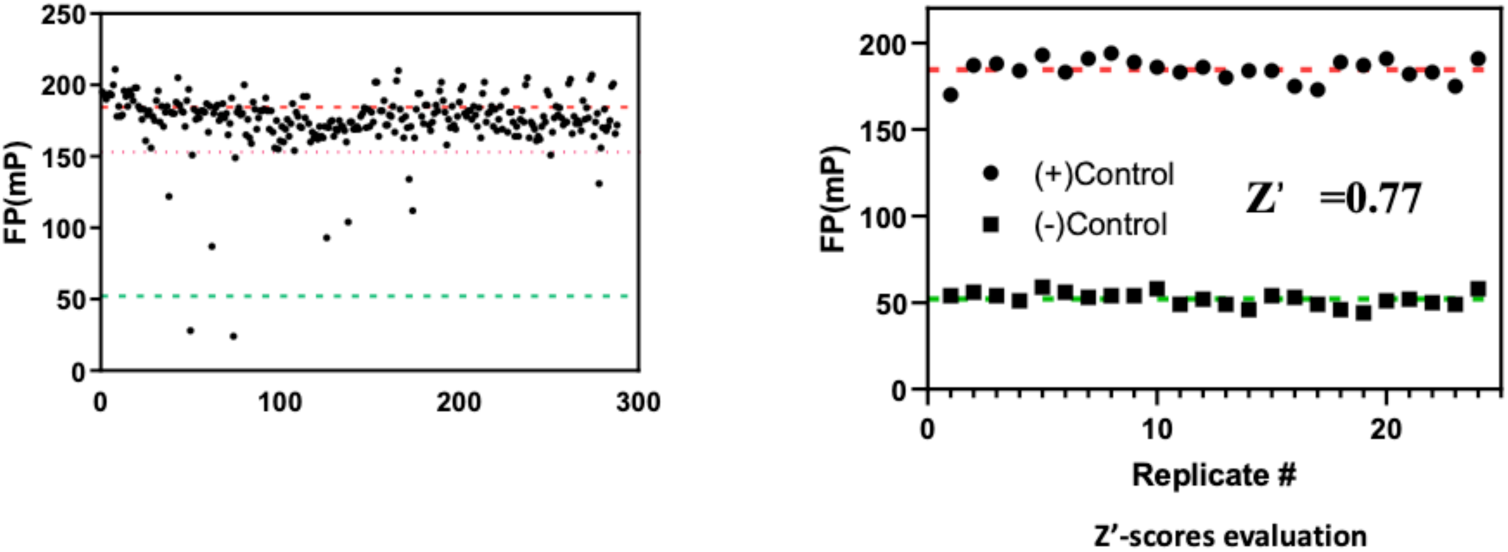
Evaluation of the FP assay in HTS formats. A) Pilot HTS of marine natural product library. B) Z’ scores evaluation.

## 4. Discussion and Conclusions

SARS-CoV-2 continual to be a big threat to public health after the initial outbreak a year ago. The membrane fusion process employed by SARS-CoV-2 to enter the host cell is a highly conserved mechanism shared by many different types of virus, suggesting that it is an attractive target to develop broad-spectrum antivirals.

Currently, research mainly relies on cell-based phenotypic assays to screen and evaluate the SARS-CoV-2 fusion inhibitors. Cell-based assays have advantages such as cell permeability factor already included, however, an economical, robust, and high-throughput *in vitro* assay is also an important complementary tool for screening and evaluation of binding potency novel fusion inhibitors, and the further structure-activity relationship studies. In this study, we developed such an *in vitro* assay to evaluate the inhibition of 6-HB by determining the binding between 5-HB and fluorescently labeled HR2P with FP techniques.

FP-based assays typically required a large amount of protein, and we solved the 5-HB production problem by fusing it to a highly soluble NusA tag, and the resulting NusA-5HB construct can be produced in large quantities in *E. coli* (30-50 mg/L). In competitive FP experiments, the IC_50_ of HR2P and EK1 peptides were determined to be 6 and 2.5 nM respectively. Five Sal family natural products were evaluated with our assay, and we found Sal A has a comparable activity with Sal C, while the other three analogs have much lower activities. Interestingly, the simple monomer, caffeic acid also has a sub-mM activity, thus could potentially serval as a starting fragment for “fragment-based drug design”.

Although our attempt to find fusion inhibitors from a small marine natural product library is not successful, the assay itself did demonstrate robustness with a Z’ factor close to 0.8 in HTS formats. We envision that this assay could be applied to screen large commercial compound libraries as the production of large quantities of NusA5HB and HR2P-FL is very straightforward. Another potential application of this assay is to screen fragment library as our assay can pick up sub-mM week activities, and the resulting fragment can be further optimized to increase the binding affinity with the aid of this assay.

## Acknowledgements

This study was supported by The National Key R&D Program of China (2019YFC0312501), Guangdong Marine Economy Promotion Fund (Grant GDOE[2019]A21). We thank Prof. Lu Lu for providing peptide EK1.

---

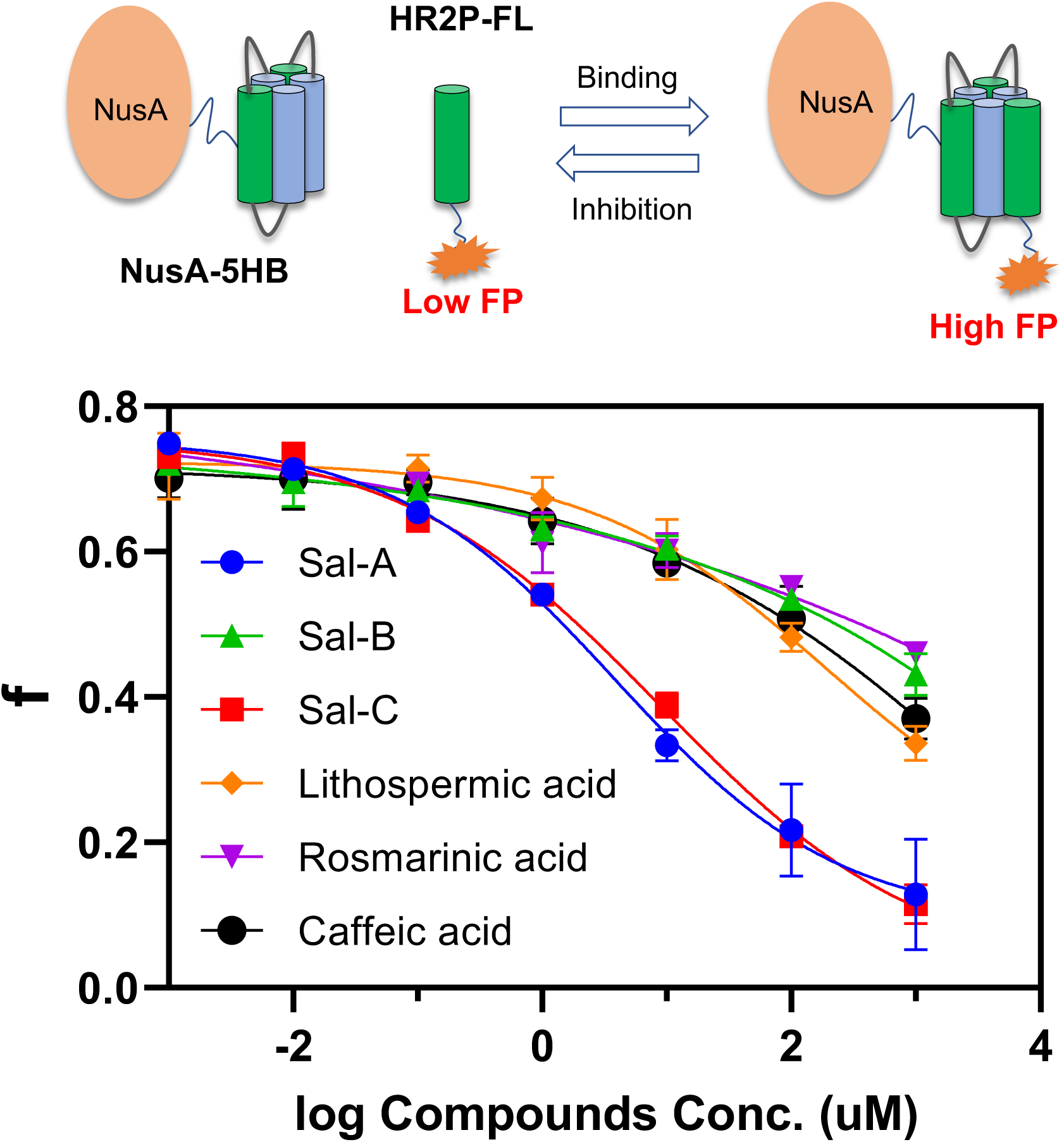

## Ref

[1] Zhou, P.; Yang, X.-L.; Wang, X.-G.; et al., A pneumonia outbreak associated with a new coronavirus of probable bat origin. nature 2020, 579, 270.

[2] of the International, C. S. G., The species Severe acute respiratory syndrome-related coronavirus: classifying 2019-nCoV and naming it SARS-CoV-2. Nature microbiology 2020, 5, 536.

[3] Sahebnasagh, A.; Avan, R.; Saghafi, F.; et al., Pharmacological treatments of COVID-19. Pharmacol Rep 2020, 72, 1446.

[4] Holshue, M. L.; DeBolt, C.; Lindquist, S.; et al., First case of 2019 novel coronavirus in the United States. New England Journal of Medicine 2020.

[5] Wrapp, D.; Wang, N.; Corbett, K. S.; et al., Cryo-EM structure of the 2019-nCoV spike in the prefusion conformation. Science 2020, 367, 1260.

[6] Xia, S.; Yan, L.; Xu, W.; et al., A pan-coronavirus fusion inhibitor targeting the HR1 domain of human coronavirus spike. Sci Adv 2019, 5, eaav4580.

[7] Martens, S.; McMahon, H. T.; Mechanisms of membrane fusion: disparate players and common principles. Nature reviews Molecular cell biology 2008, 9, 543.

[8] Podbilewicz, B.; Virus and cell fusion mechanisms. Annual review of cell and developmental biology 2014, 30, 111.

[9] Matthews, T.; Salgo, M.; Greenberg, M.; et al., Enfuvirtide: the first therapy to inhibit the entry of HIV-1 into host CD4 lymphocytes. Nature reviews Drug discovery 2004, 3, 215.

[10] Outlaw, V. K.; Bovier, F. T.; Mears, M. C.; et al., Inhibition of coronavirus entry in vitro and ex vivo by a lipid-conjugated peptide derived from the sars-cov-2 spike glycoprotein hrc domain. Mbio 2020, 11.

[11] Xia, S.; Liu, M.; Wang, C.; et al., Inhibition of SARS-CoV-2 (previously 2019-nCoV) infection by a highly potent pan-coronavirus fusion inhibitor targeting its spike protein that harbors a high capacity to mediate membrane fusion. Cell Res 2020, 30, 343.

[12] Xia, S.; Zhu, Y.; Liu, M.; et al., Fusion mechanism of 2019-nCoV and fusion inhibitors targeting HR1 domain in spike protein. Cell Mol Immunol 2020, 17, 765.

[13] Zhu, Y.; Yu, D.; Yan, H.; et al., Design of potent membrane fusion inhibitors against SARS-CoV-2, an emerging coronavirus with high fusogenic activity. Journal of virology 2020, 94.

[14] Wang, X.; Xia, S.; Wang, Q.; et al., Broad-spectrum coronavirus fusion inhibitors to combat COVID-19 and other emerging coronavirus diseases. International journal of molecular sciences 2020, 21, 3843.

[15] Lu, H.; Zhou, Q.; He, J.; et al., Recent advances in the development of protein-protein interactions modulators: mechanisms and clinical trials. Signal Transduct Target Ther 2020, 5, 213.

[16] Atanasov, A. G.; Zotchev, S. B.; Dirsch, V. M.; et al., Natural products in drug discovery: advances and opportunities. Nature Reviews Drug Discovery 2021, 1.

[17] Yang, C.; Pan, X.; Xu, X.; et al., Salvianolic acid C potently inhibits SARS-CoV-2 infection by blocking the formation of six-helix bundle core of spike protein. Signal transduction and targeted therapy 2020, 5, 1.

[18] Hoffer, L.; Voitovich, Y. V.; Raux, B.; et al., Integrated strategy for lead optimization based on fragment growing: the diversity-oriented-target-focused-synthesis approach. Journal of medicinal chemistry 2018, 61, 5719.

[19] Hall, M. D.; Yasgar, A.; Peryea, T.; et al., Fluorescence polarization assays in high-throughput screening and drug discovery: a review. Methods and applications in fluorescence 2016, 4, 022001.

[20] Roehrl, M. H.; Wang, J. Y.; Wagner, G.; A general framework for development and data analysis of competitive high-throughput screens for small-molecule inhibitors of protein− protein interactions by fluorescence polarization. Biochemistry 2004, 43, 16056.

[21] Zhang, J.-H.; Chung, T. D.; Oldenburg, K. R.; A simple statistical parameter for use in evaluation and validation of high throughput screening assays. Journal of biomolecular screening 1999, 4, 67.

[22] Uri, A.; Nonga, O. E.; What is the current value of fluorescence polarization assays in small molecule screening? Expert opinion on drug discovery 2020, 15, 131.

[23] Root, M. J.; Kay, M. S.; Kim, P. S.; Protein design of an HIV-1 entry inhibitor. Science 2001, 291, 884.

[24] Frey, G.; Rits-Volloch, S.; Zhang, X.-Q.; et al., Small molecules that bind the inner core of gp41 and inhibit HIV envelope-mediated fusion. Proceedings of the National Academy of Sciences 2006, 103, 13938.

[25] De Marco, V.; Stier, G.; Blandin, S.; et al., The solubility and stability of recombinant proteins are increased by their fusion to NusA. Biochemical and biophysical research communications 2004, 322, 766.

[26] De Marco, A.; Two-step metal affinity purification of double-tagged (NusA–His 6) fusion proteins. Nature protocols 2006, 1, 1538.

[27] Wang, J.; Pan, X.; Tien, P.; et al., High level expression of 5-helix protein in HIV gp41 heptad repeat regions and its virus fusion-inhibiting activity. Sheng wu gong cheng xue bao= Chinese journal of biotechnology 2009, 25, 435.

[28] Jameson, D. M.; Ross, J. A.; Fluorescence polarization/anisotropy in diagnostics and imaging. Chemical reviews 2010, 110, 2685.

[29] Irving, M.; Steady-state polarization from cylindrically symmetric fluorophores undergoing rapid restricted motion. Biophysical journal 1996, 70, 1830.

[30] Huang, X.; Fluorescence polarization competition assay: the range of resolvable inhibitor potency is limited by the affinity of the fluorescent ligand. Journal of biomolecular screening 2003, 8, 34.

[31] Gao, Z.; Ovchinnikova, O. G.; Huang, B.-S.; et al., High-throughput “FP-tag” assay for the identification of glycosyltransferase inhibitors. Journal of the American Chemical Society 2019, 141, 2201.

[32] Yin, X.; Li, J.; Chen, S.; et al., An Economical High-Throughput “FP-Tag” Assay for Screening Glycosyltransferase Inhibitors. ChemBioChem 2020.

