## Supplemental Fig for "A Robust High-throughput Fluorescent Polarization Assay for the Evaluation and Screening of SARS-CoV-2 Fusion Inhibitors"

[b] Prof. Lan Liu, Prof. Zhizeng Gao

### Contents List

**Figure S1.** SDS-PAGE analysis of the purified proteins

**Figure S2.** Structure of NusA-5HB gene

**Table S1.** Gene sequences of NusA-5HB and 5HB

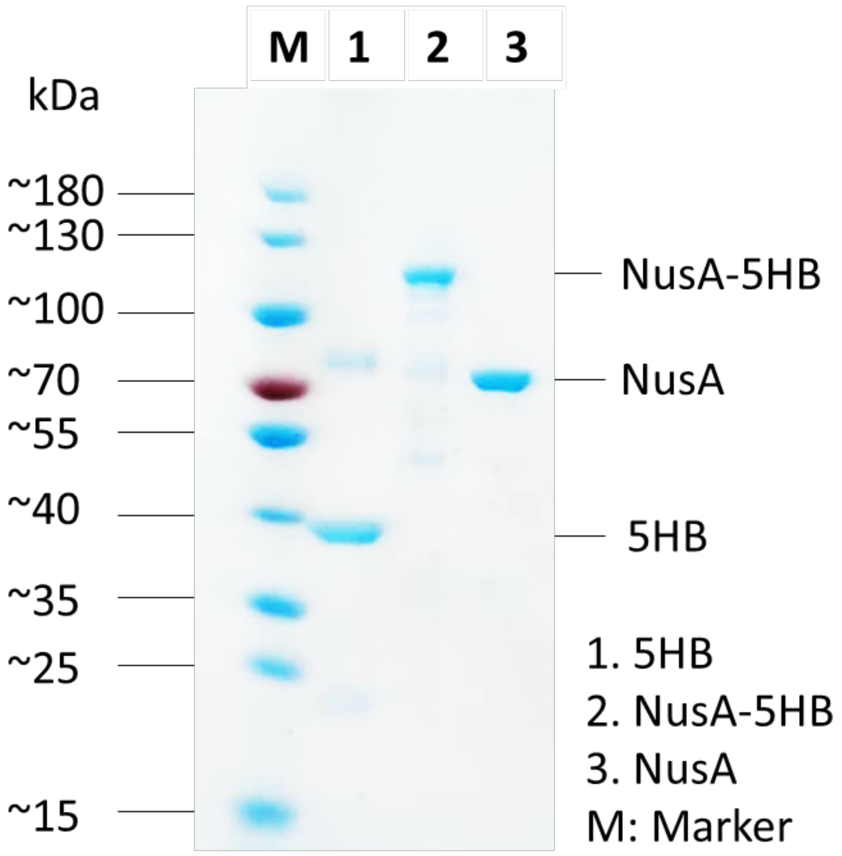


**Figure S1**. **SDS-PAGE analysis of the purified proteins**. Lane M: molecular weight marker, Lane 1: purified 5HB, Lane 2: purified NusA-5HB, Lane 3: purified NusA.

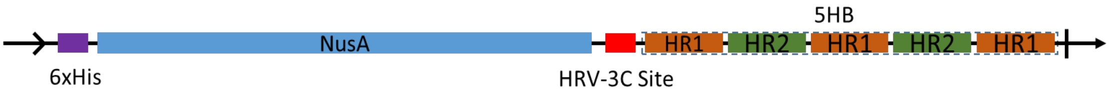


**Figure S2. General structure of NusA-5HB gene**
